# MAS Cryoprobe Enhances Solid-State NMR Signals of α-Synuclein Fibrils

**DOI:** 10.1101/2025.09.20.677435

**Authors:** Malitha C. Dickwella Widanage, Barbara Perrone, Jhinuk Saha, Riqiang Fu, Jochem Struppe, Ryan P. McGlinchey, Jennifer C. Lee, Robert W. Schurko, Ayyalusamy Ramamoorthy

## Abstract

Solid-state NMR spectroscopy is increasingly applied to structural and dynamics studies across a broad range of chemical, material, and biological systems. Although sensitivity has traditionally been a major limitation, the recently developed MAS cryoprobe has been shown to substantially overcome this challenge. Its ability to enhance the signal-to-noise (S/N) ratio without requiring sample freezing makes it particularly attractive for investigating non-isotropic systems, including soft materials (e.g., hydrogels), semi-solids (e.g., membrane mimetics) and rigid solids (e.g., amyloid fibrils). In this study, we report on the enhanced sensitivity of solid-state NMR experiments on α-synuclein fibrils using a MAS cryoprobe. Nearly an order-of-magnitude improvement in S/N was observed in CPMAS, refocused-INEPT and 2D ^13^C-^13^C chemical shift correlation spectra of α-synuclein fibrils compared with data collected on a conventional MAS probe. The improved S/N enables the acquisition of slowly decaying signals in the indirect dimension, facilitating faster, high-resolution multidimensional solid-state NMR spectroscopy. We therefore anticipate that MAS cryoprobe will become increasingly valuable for structural studies a wide range of samples that are less abundant, less stable, or transient, such as amyloid intermediates.

## Introduction

Solid-state NMR spectroscopy is widely used for atomistic-resolution structural and dynamics studies of a variety of solids. Magic-angle spinning (MAS) combined with recoupling techniques has become a standard approach for characterizing the structure and dynamics of amyloid fibrils.[1-7] While many high-resolution structures of amyloid fibrils have been reported, the investigation of lowly populated or heterogeneous amyloid species remains challenging due to the inherently low sensitivity of solid-state NMR spectroscopy.

Although dynamic nuclear polarization (DNP) can substantially enhance sensitivity,[8-12] room-temperature measurements are often preferred for studying dynamic species and preserving native-like sample conditions. This is especially important for membrane-associated proteins; membrane-active biomolecules such as antimicrobial peptides, toxins, amyloid peptides, and fusion peptides; transient intermediates such as amyloid oligomers; and other dynamic systems. In this context, the recently developed MAS cryoprobe offers a valuable alternative by significantly enhancing the sensitivity and without the need to freeze the sample.[13-18]

In this study, we report solid-state NMR spectroscopy results acquired using a MAS cryoprobe on uniformly ^13^C- and ^15^N-labeled α-synuclein (α-Syn) fibrils formed in the presence of 500 mM NaCl. α-Syn fibril deposition is a key pathological feature of neurodegenerative disorders such as Parkinson’s disease, dementia with Lewy bodies, and multiple system atrophy.[19, 20] Importantly, α-syn fibrils are classified as amyloids, adopting cross-β structure, stabilized by hydrogen-bond networks of amide backbones, steric zippers, and salt bridges, as determined by solid-state NMR [21, 22] and other high-resolution techniques.[23-28] Fibril structural characterization is critical to understanding disease mechanisms as recent studies indicate a possible relationship between distinct fibril polymorphs and disease phenotypes.[21, 29-31] Since α-syn fibrils are highly polymorphic and concentrations of these structural variants differ within a sample, it becomes challenging to study them by solid-state NMR spectroscopy. Spectral line-broadening hampers resolution, and more critically, low-abundance amyloid species remain undetected. However, solid-state NMR experiments performed using a MAS cryoprobe, without the need to freeze the sample, have been shown to enhance sensitivity, thereby potentially enabling the detection of minor amyloid species in heterogeneous mixtures.

## Materials and Methods

### Fibril preparation

Expression and purification of uniformly labeled α-synuclein was carried as described.[32] Monomeric α-synuclein at a final concentration of 150 μM was incubated in 10 mM sodium phosphate buffer (pH 7.4) containing 500 mM NaCl (∼1 mL total volume), under continuous agitation at 700 rpm in a benchtop thermal mixer (Fisherbrand) at 37 °C for 5 days to promote fibril formation. After incubation, the samples were centrifuged at 17,000 rpm for 30 minutes at 4 °C to pellet the fibrils, and the supernatant containing soluble oligomers or monomers was carefully discarded. Approximately ∼1 mg of the fibrillar pellet was collected and lyophilized to remove residual water. The dried fibrils were then resuspended in 30-35 μL of D_2_O (with a final buffer concentration of Tris at 3.5 mM, NaPi at 1.42 mM, and NaCl at 75 mM, and 2 mM of α-synuclein) and tightly packed in a 3.2 mm MAS NMR rotor (Bruker Cryo-MAS probe and a home-built probe at the MagLab) for solid-state NMR measurements.

### NMR experiments

All experiments were performed on Bruker Avance 600 MHz (14.1 T) spectrometers equipped with a 3.2 mm HCN MAS Cryoprobe (Bruker) and a conventional-MAS (home-built at the MagLab (NHMFL)) probe operating at 12 kHz MAS and a regulated temperature of 268 K. ^13^C chemical shifts were externally referenced to adamantane (CH_2_ resonance at 38.48 ppm, relative to TMS at 0 ppm), and ^15^N shifts were referenced to the methionine amide nitrogen (at 127.88 ppm) of N-formyl-Met-Leu-Phe-OH on the liquid ammonia scale (at 0 ppm). Typical RF field strengths used were 62.5 kHz for ^1^H decoupling, 62.5–88 kHz for ^1^H hard pulses, and 45–60 kHz for ^13^C. All NMR experimental parameters and spectral processing parameters used in this study are given in Table 1.

**Table 1.** A summary of experimental and data processing parameters used to carry out solid-state NMR experiments reported in this study.

| Experiment | NMR parameters |  |  |  |  |  |  |  |  |  |  | Processing parameters |
| --- | --- | --- | --- | --- | --- | --- | --- | --- | --- | --- | --- | --- |
|  | T (K) | B<sub>0</sub> (T) | v<sub>MAS</sub> (kHz) | ns | d1 (s) | t<sub>1, max</sub> (ms) | t<sub>1, inc</sub> (μs) | τ<sub>dw</sub> (μs) | τ<sub>acq</sub> (ms) | τ<sub>HC</sub> (ms) | τ<sub>SD</sub> (ms) |  |
| MAS Cryoprobe |  |  |  |  |  |  |  |  |  |  |  |  |
| 1D <sup>13</sup>C CP | 268 | 14.1 | 12 | 128 | 2 |  |  | 11 | 23 | 1 |  | EM (LB: 25 Hz) |
| 1D <sup>15</sup>N CP | 268 | 14.1 | 12 | 1024 | 2 |  |  | 21 | 43 | 1 |  | EM (LB: 10 Hz) |
| 1D <sup>13</sup>C Refocused-INEPT | 268 | 14.1 | 12 | 1024 | 2 |  |  | 11 | 49 |  |  | EM (LB: 50 Hz) |
| 2D <sup>13</sup>C-<sup>13</sup>C DARR | 268 | 14.1 | 12 | 2<br>-32 | 2<br>-3 | 18 | 33 | 5 | 23 | 1 | 50-250 | QSINE (SSB: 4) |
| Conventional MAS probe |  |  |  |  |  |  |  |  |  |  |  |  |
| 1D <sup>13</sup>C CP | 268 | 14.1 | 12 | 128 | 2 |  |  | 5 | 10 | 1 |  | EM (LB: 25 Hz) |
| 1D <sup>15</sup>N CP | 268 | 14.1 | 12 | 1024 | 3 |  |  | 20 | 20 | 1.5 |  | EM (LB: 10 Hz) |
| 1D <sup>13</sup>C Refocused-INEPT | 268 | 14.1 | 12 | 1024 | 2 |  |  | 5 | 20 |  |  | EM (LB: 50 Hz) |
| 2D <sup>13</sup>C-<sup>13</sup>C DARR | 268 | 14.1 | 12 | 32 | 2 | 10 | 20 | 5 | 10 | 1 | 50-250 | QSINE (SSB: 4) |

T, sample temperature; B<sub>0</sub>, magnetic field; v<sub>MAS</sub>, MAS frequency; ns, number of scans; d<sub>1</sub>, recycle delay or delay between successive scans; t<sub>1,max</sub>, maximum t<sub>1</sub> evolution time (for the indirect dimension); t<sub>1,inc</sub>, time increment for t<sub>1</sub> (for the indirect dimension) evolution time; τ<sub>dw</sub>, dwell time used in the direct dimension; τ<sub>acq</sub>, maximum acquisition time; τ<sub>XY</sub>, cross-polarization contact time for <sup>1</sup>H to <sup>13</sup>C or <sup>15</sup>N; τ<sub>SD</sub>, DARR (Dipolar-Assisted Rotational Resonance) mixing or spin diffusion time.

One-dimensional ^13^C NMR spectra were acquired using selective polarization transfer schemes to distinguish between the rigid and dynamic components of α-synuclein fibrils. Rigid, immobilized domains were probed using ^13^C cross-polarization (CP) MAS with a contact time of 1 ms, optimized to enhance magnetization transfer from ^1^H to ^13^C via dipolar couplings that are preserved in the solid-state. To selectively detect mobile and conformationally flexible regions, particularly side chains and disordered termini, refocused-INEPT (Insensitive Nuclei Enhanced by Polarization Transfer) experiments were employed. This through-bond magnetization transfer sequence utilizes scalar J-couplings (J_CH_) between ^1^H and ^13^C nuclei. The refocused-INEPT block included two delay elements of 1/4J and 1/6J (calculated using a coupling constant of 170 Hz, typical for a CH group), allowing efficient polarization transfer from protons to carbons in highly
dynamic environments. Given the rapid transverse relaxation of rigid components, only segments with high internal mobility exhibiting long T_2_ (spin-spin relaxation) times, primarily dynamic side chains, contribute to the INEPT-detected signals. Two-dimensional (2D) ^13^C-^13^C chemical shift correlation spectra were acquired using the Dipolar-Assisted Rotational Resonance (DARR) technique with 50 and 250 ms mixing times.[33] This experiment enables homonuclear magnetization transfer via dipolar couplings under MAS, facilitating the detection of intramolecular ^13^C-^13^C cross-peaks, particularly between Cα, Cβ, and side chain carbons of structurally informative residues. DARR spectra provides insights into local secondary structure and residue-specific packing within the fibril core.

## Results and Discussion

In this study, we compare solid-state NMR spectra obtained using a MAS cryoprobe with those acquired using a conventional MAS probe under identical experimental conditions. The performance of a 3.2 mm HCN MAS cryoprobe (from Bruker) was benchmarked against the conventional 3.2 mm HCN MAS probe (home-built MagLab) for solid-state NMR of ∼1 mg α-synuclein fibrils. ^13^C and ^15^N-labeled α-synuclein was expressed and purified as reported elsewhere,[32] and the fibril samples were prepared as described in the Supporting Information. 1D ^13^C and ^15^N cross-polarization (CP) experiments were performed at under 12 kHz MAS at 268 K on a 600 MHz (14.1 T) NMR spectrometer (**Fig. 1**). The use of MAS cryoprobe provided significantly enhanced sensitivity, allowing the acquisition of high-quality two-dimensional spectra with a reduced experimental time as explained below.

As shown in **Fig. 1**, the data collected with the MAS cryoprobe (yellow traces) exhibit pronounced sensitivity improvements for both ^13^C and ^15^N nuclei. The ^13^C MAS spectra (**Fig. 1a**) show that weak aliphatic (∼65 to 75 ppm) and aromatic (∼105 to 135 ppm) resonances are better resolved in the spectrum acquired using the MAS cryoprobe. Quantitative signal-to-noise (S/N) analysis yields a sensitivity gain of about 6 for ^13^C, which is in excellent agreement with previously reported gains for similar experiments using a MAS cryoprobe.[28] Similarly, the ^15^N CP spectra (**Fig. 1b**) demonstrate a remarkable ∼9-fold S/N enhancement when using a MAS cryoprobe, facilitating the detection of amide signals that are barely discernible in the spectrum acquired using the conventional probe. The improved sensitivity reflects the reduced thermal noise and the enhanced cryogenic preamplifier efficiency of the MAS cryoprobe design, which is particularly advantageous for low-γ nuclei like ^15^N and for samples available in limited quantities.

**Figure 1.**
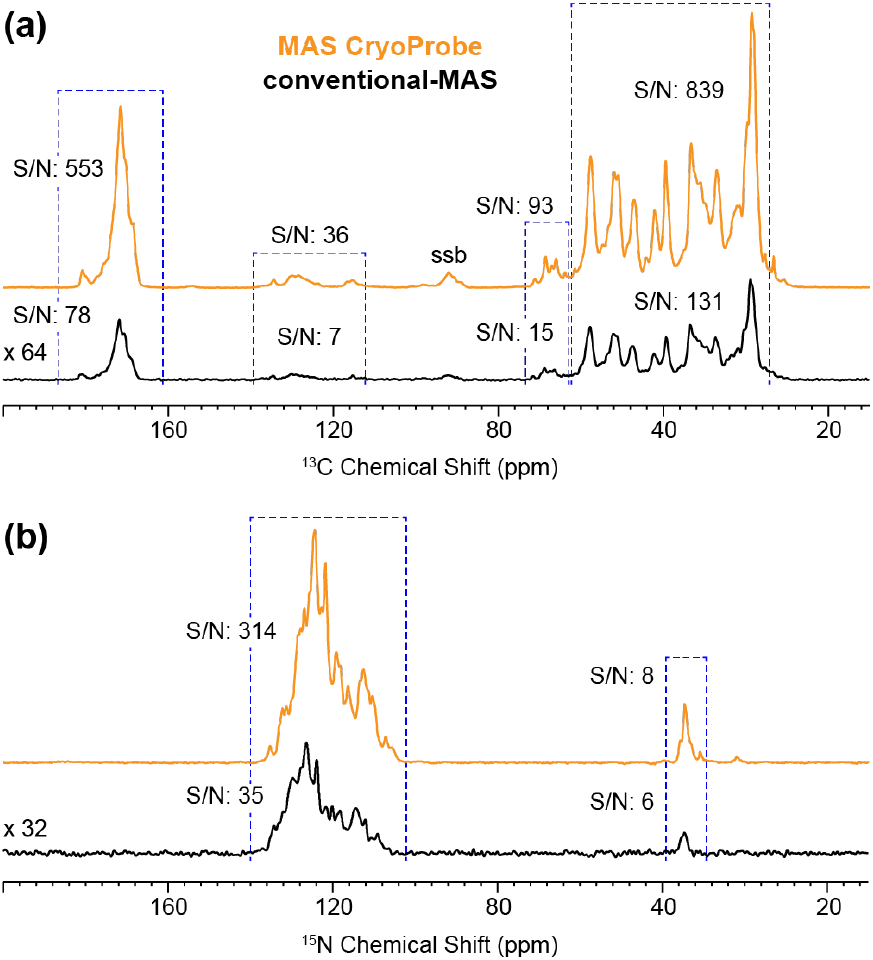
MAS cryoprobe enhances the sensitivity for both ^13^C- and ^15^N-detection. (a) ^13^C CPMAS spectra and (b) ^15^N CPMAS spectra acquired using a MAS cryoprobe (Bruker, yellow) and a conventional 3.2 mm HCN MAS probe (home-built at the MagLab, black). Measurements were performed on a 600 MHz spectrometer at 12 kHz MAS and 268 K with ∼1 mg of α-synuclein fibrils. All NMR parameters are given in Table 1.

The MAS cryoprobe provided pronounced enhancements in both signal sensitivity and spectral resolution for 2D ^13^C-^13^C correlation experiments. As illustrated in **Fig. 2a**, spectra acquired with a 50 ms DARR mixing time using the MAS cryoprobe demonstrate substantially higher S/N ratio relative to those obtained using a conventional MAS probe under identical acquisition parameters (NS = 32). Remarkably, even with minimal signal averaging (NS = 2), the MAS cryoprobe spectrum retains comparable, or even better, spectral details, emphasizing the significant reduction in experimental time afforded by MAS cryoprobe (**Fig. 2b**). The application of a QSINE apodization function during Fourier transformation further improved spectral quality by attenuating truncation artifacts while preserving intrinsic resolution.[15, 34]

**Figure 2.**
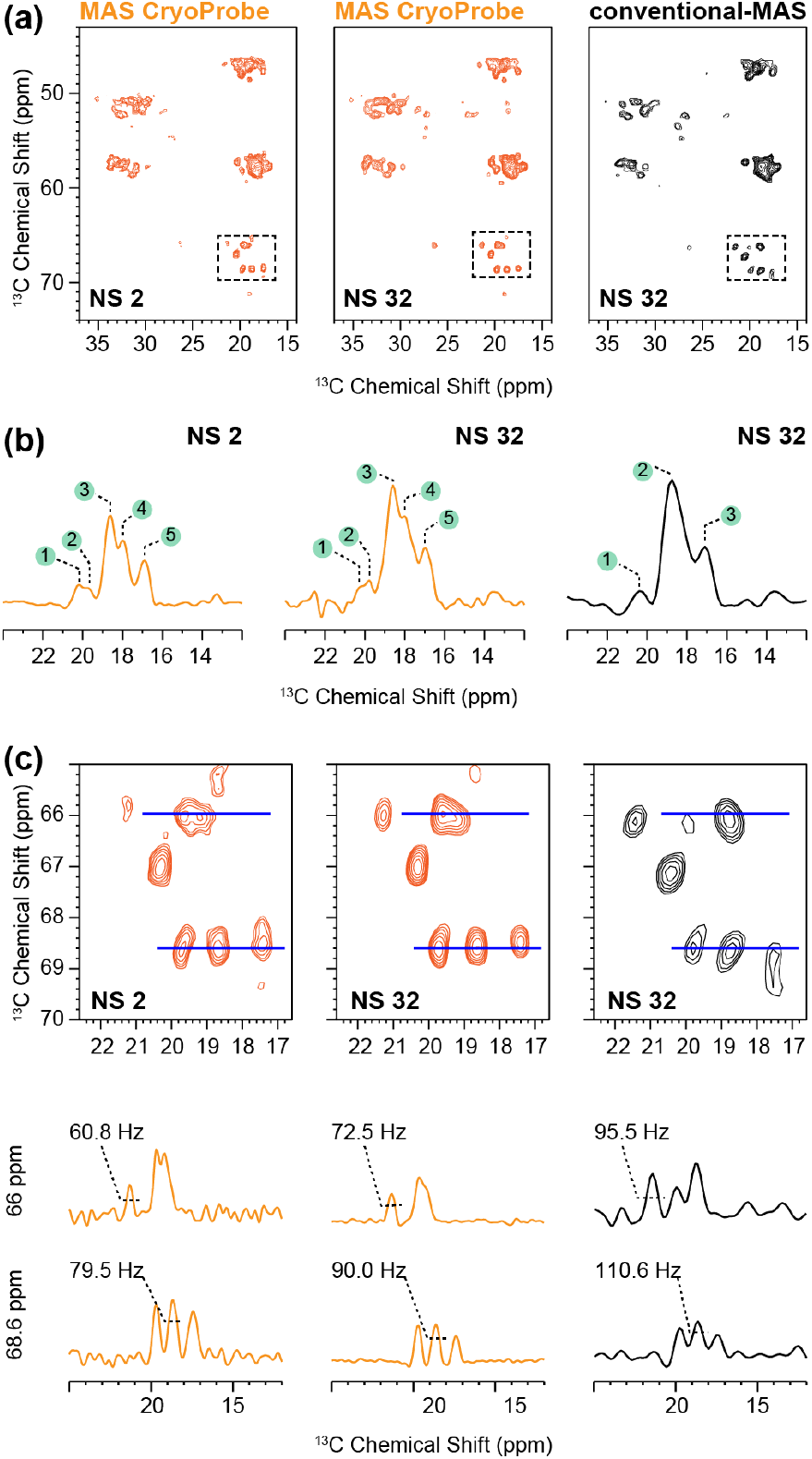
Signal enhancement and improved resolution with MAS cryoprobe. 2D ^13^C-^13^C spectra of α-synuclein fibrils acquired with 50 ms DARR (a) using a MAS cryoprobe (left: number of accumulated scans (NS) = 2; middle: NS = 32) and a conventional MAS probe (right: NS = 32), processed with a QSINE apodization function. Selected 1D slices (b) at 57.8 ppm demonstrate the enhanced sensitivity and resolution achieved with MAS cryoprobe. Expanded views (c) of 2D spectral regions (highlighted with dashed line box), with blue lines indicating the positions of additional 1D slices at 66.0 ppm and 68.6 ppm. Full width at half maximum (FWHM) values are provided for selected peaks to illustrate resolution differences. All the NMR and processing parameters used to obtain these spectra are given in Table 1.

To further demonstrate the sensitivity and resolution gains obtained with MAS cryoprobe, selected 1D spectral slices extracted from 2D spectra are shown in **Fig. 2b**. Notably, the enhanced resolution enables the discrimination of five distinct peaks, whereas the conventional MAS probe yields only three broadened and unresolved resonances (14– 22 ppm). Expanded views of the highlighted spectral regions (**Fig. 2c**) further reveal narrow linewidths in spectra acquired with the MAS cryoprobe, as quantified by full width at half maximum (FWHM) values for peaks at 66.0 ppm and 68.6 ppm. While a cryoprobe does not inherently provide higher spectral resolution, the improved resolution observed in Fig.2 is a consequence of the enhanced S/N.

Employing a MAS cryoprobe yields a marked enhancement in spectral sensitivity and retention of long-range ^13^C-^13^C correlations in 2D DARR experiments at extended mixing times. As shown in **Fig. 3a**, the 250 ms DARR spectrum acquired with a MAS cryoprobe exhibits substantially higher cross-peak intensities and S/N compared to that obtained with the conventional MAS probe, reflecting the improved detection sensitivity afforded by the cryoprobe.[35] The ability to detect weak dipolar coupling driven magnetization transfers during long mixing periods enables the observation of inter-residue and distal intra-residue correlations. Extracted 1D traces along the direct (t2) and indirect (t1) dimensions (**Fig. 3b** and **3c**, respectively) highlight the enhanced dynamic range and resolution achievable due to the enhanced S/N rendered by a MAS cryoprobe. This sensitivity gain would be valuable for the complete mapping of structural connectivity in complex solid-state systems, particularly for aliphatic and aromatic spin systems where cross-peak intensities are inherently low.

**Figure 3.**
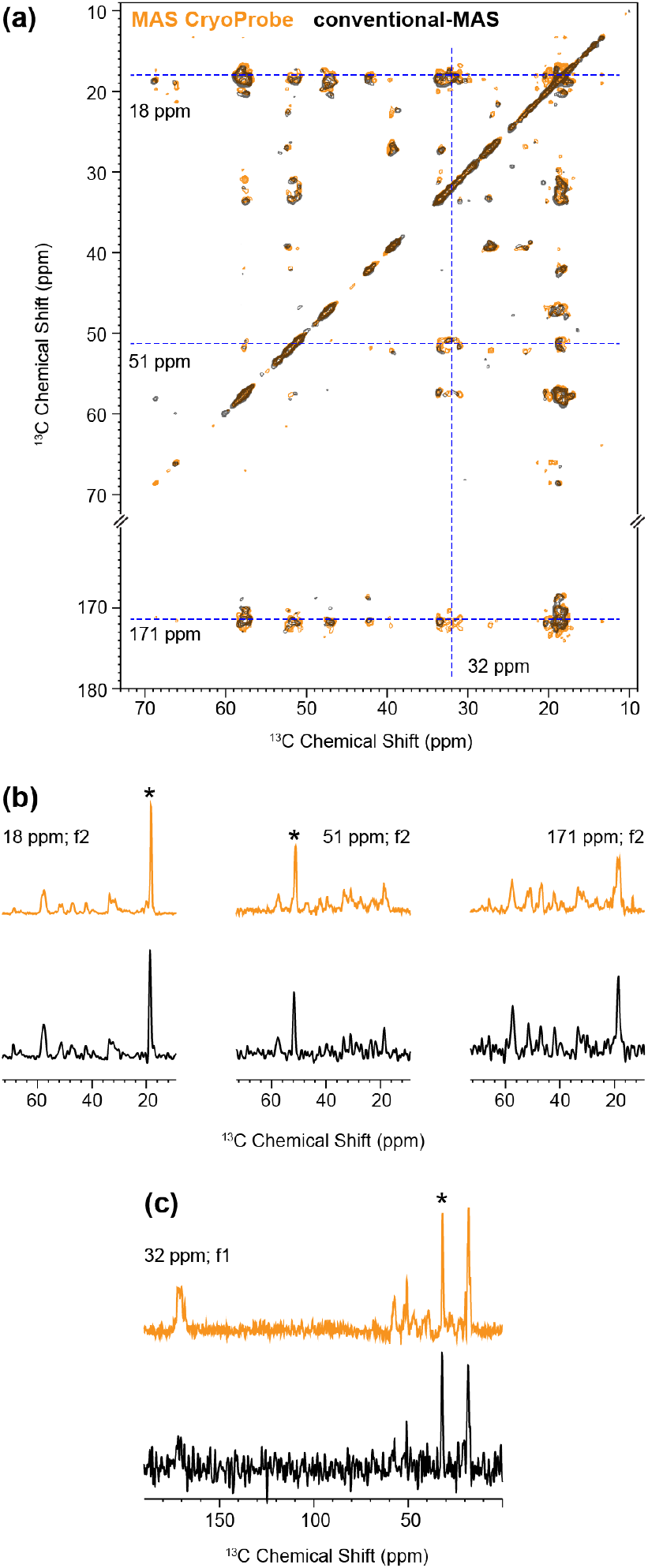
Long mixing time preserves signal intensity with MAS cryoprobe. 2D ^13^C-^13^C 250 ms DARR spectra of α-synuclein fibrils acquired using MAS cryoprobe and, conventional MAS probes (a). (b and c) Selected 1D slices extracted at the frequencies marked by blue dashed lines are shown for the direct (t2) and the indirect (t1) dimensions. Spectra illustrate enhanced cross-peak retention and sensitivity in MAS cryoprobe. All the NMR and processing parameters are given in Table 1. * denotes the diagonal peak.

The ^13^C refocused-INEPT spectra acquired with the MAS cryoprobe also exhibit a pronounced sensitivity enhancement, particularly in the aliphatic region, compared to that obtained using a conventional MAS probe (**Fig. 4**). Refocused-INEPT selectively can be used to selectively detect mobile molecular components (or regions of a protein), by using scalar (J) coupling-based coherence transfer that is efficient only in segments undergoing fast motion. Consequently, it emphasizes flexible aliphatic chains, dynamic side groups, and mobile carbohydrate or lipid moieties whose signals are typically attenuated in CP-based experiments due to weak ^1^H-^13^C dipolar coupling. The first FID obtained from the 2D refocused-INEPT based 2D ^1^H/^13^C HETCOR experiment acquired with MAS cryoprobe (bottom) exhibits higher S/N, further illustrating the enhanced performance of MAS cryoprobe for the detection of dynamic components in the sample (**Supplementary Fig. 1**).

**Figure 4.**
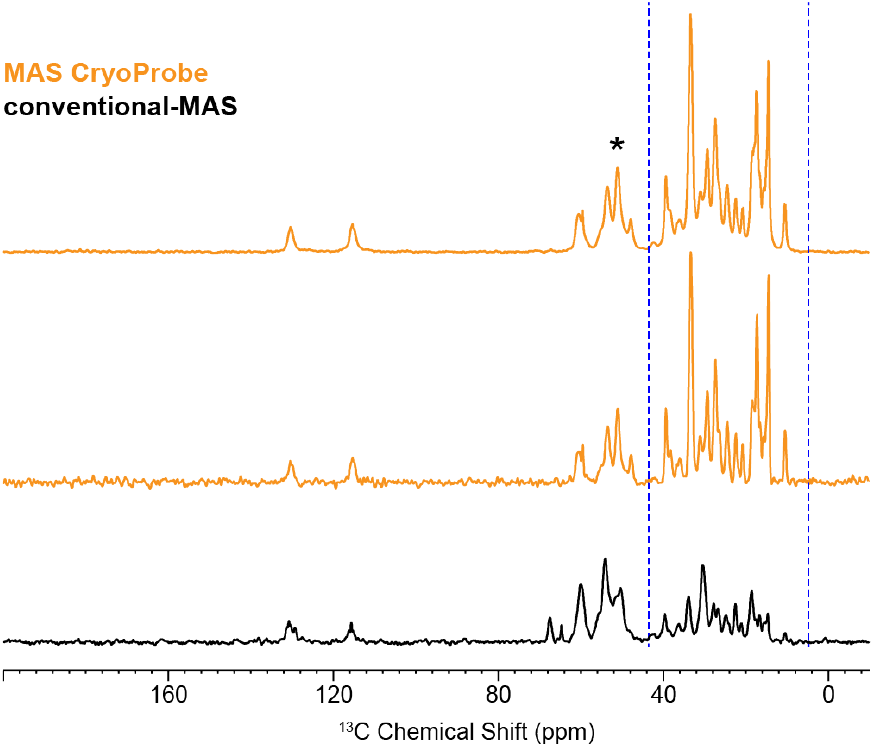
Enhanced sensitivity in the aliphatic region with MAS cryoprobe. 1D ^13^C refocused-INEPT spectra of α-synuclein fibrils acquired using MAS cryoprobe (top) along with the 1st FID of the 2D ^1^H/^13^C HETCOR from MAS cryoprobe (middle), and 1D ^13^C refocused-INEPT spectrum acquired using conventional MAS probe (bottom). All spectra were locally normalized to the region indicated by asterisks to highlight relative signal enhancement. All the NMR parameters used to acquire these spectra are given in Table 1.

## Conclusions

Here, we reported solid-state NMR experiments on α-synuclein amyloid fibrils carried out under MAS using a cryoprobe. The sensitivity gains achieved using the MAS cryoprobe are compared with those obtained using a conventional MAS probe. The reported experimental results demonstrate the advantage of employing a cryogenic probe for solid-state NMR studies of biological solids, particularly in cases where the sample quantity is limited or the desired nucleus has an inherently low receptivity, such as ^15^N. The increased sensitivity directly translates into reduced experimental time or improved spectral quality through additional signal averaging. Moreover, because slowly decaying signals often exhibit lower S/N and can limit the number of data points collected in the indirect dimension, the cryogenic probe is expected to be especially valuable for acquiring high-resolution multidimensional spectra.

## Acknowledgements

This study was funded by the NIH (R01DK132214 to A.R.). The work performed at NHMFL is supported by National Science Foundation Cooperative Agreement No. DMR-2128556* and the State of Florida. This research was supported in part by the Intramural Research Program of the National Institutes of Health (NIH). The contributions of the NIH authors are considered Works of the United States Government. The findings and conclusions presented in this paper are those of the authors and do not necessarily reflect the views of the NIH or the U.S. Department of Health and Human Services.

## Data availability

Materials and methods, supplementary figures and additional experimental data are included in the Supporting Information. Experimental data files are available at the *Open Science Framework*:https://osf.io/3k9v8/?view_only=962b2a565973422eae64bfa2463165df.

