## Supporting Information for "MAS Cryoprobe Enhances Solid-State NMR Signals of α-Synuclein Fibrils"

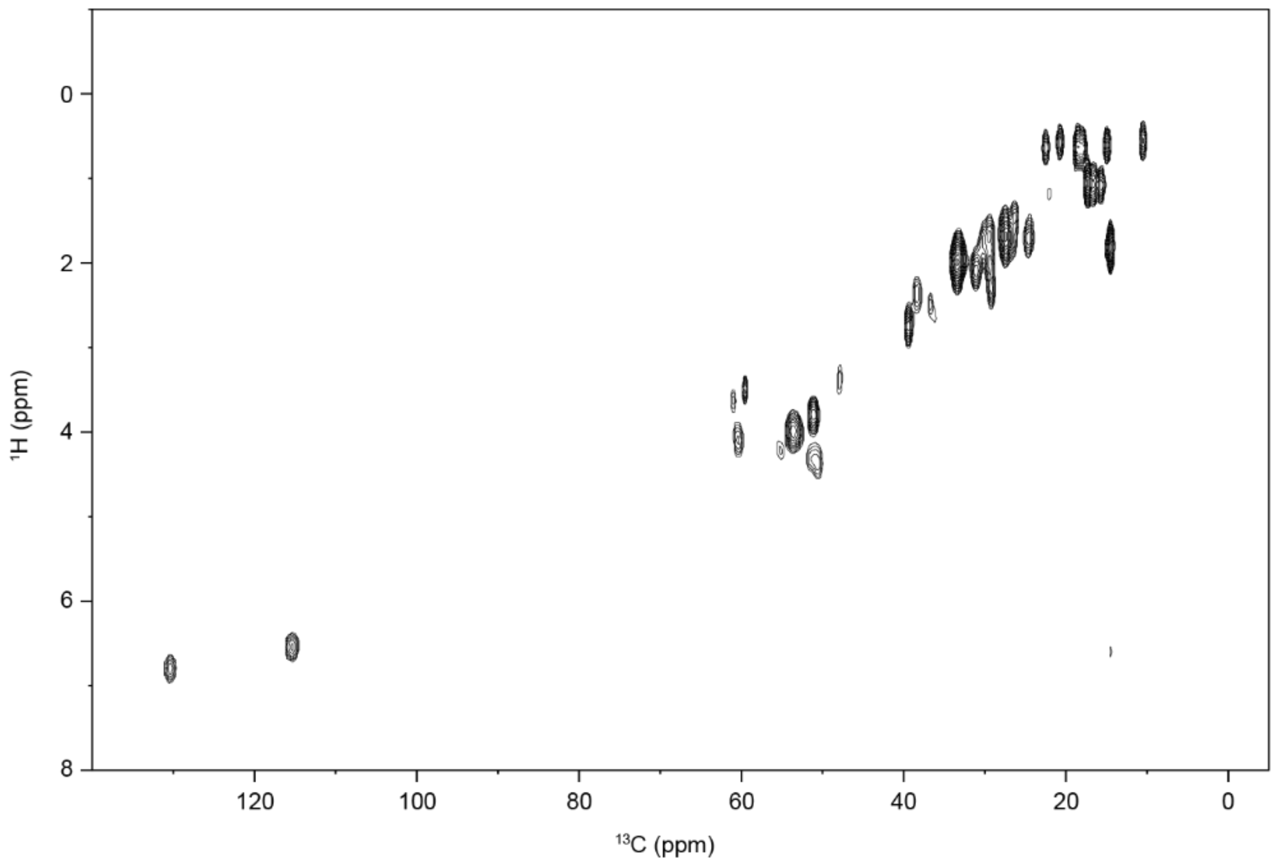


**Supplementary Figure 1. 2D $^{1}$H/$^{13}$C chemical shift correlation spectrum of α-synuclein fibrils.** Spectrum was acquired on a 600 MHz Bruker NMR spectrometer at 12 kHz MAS.
